# Predicting JNK1 Inhibitors Regulating Autophagy in Cancer using Random Forest Classifier

**DOI:** 10.1101/459669

**Authors:** Chetna Kumari, Naidu Subbarao, Muhammad Abulaish

## Abstract

Autophagy (in Greek: self-eating) is the cellular process for delivery of heterogenic intracellular material to lysosomal digestion. Protein kinases are integral to the autophagy process, and when dysregulated or mutated cause several human diseases. Atg1, the first autophagy-related protein identified is a serine/threonine protein kinases (STPKs). mTOR (mammalian Target of Rapamycin), AMPK (AMP-activated protein kinase), Akt, MAPK (mitogen-activated protein kinase) and PKC (protein kinase C) are other STPKs which regulate various components/steps of autophagy, and are often deregulated in cancer. MAPK have three subfamilies – ERKs, p38, and JNKs. JNKs (c-Jun N-terminal Kinases) have three isoforms in mammals – JNK1, JNK2, and JNK3, each with distinct cellular locations and functions. JNK1 plays role in starvation induced activation of autophagy, and the context-specific role of autophagy in tumorigenesis establish JNK1 a challenging anticancer drug target. Since JNKs are closely related to other members of MAPK family (p38, MAP kinase and the ERKs), it is difficult to design JNK-selective inhibitors. Designing JNK isoform-selective inhibitors are even more challenging as the ATP-binding sites among all JNKs are highly conserved. Although limited informations are available to explore computational approaches to predict JNK1 inhibitors, it seems diificult to find literature exploring machine learning techniques to predict JNKs inhibitors. This study aims to apply machine learning to predict JNK1 inhibitors regulating autophagy in cancer using Random Forest (RF). Here, RF algorithm is used for two purposes‐ to select and rank the molecular descriptors calculated using PaDEL descriptor software and as clasifier. The descriptors are prioritized by calculating Variable Importance Measures (VIMs) using functions based on *mean square error* (IncMSE) and *node purity* (IncNodePurity) of RF. The classification models based on a set of 22 prioritized descriptors shows accuracy 86.36%, precision 88.27% and AUC (Area Under ROC curve) 0.8914. We conclude that machine learning-based compound classification using Random Forest is one of the ligand-based approach that can be opted for virtual screening of large compound library of JNK1 bioactives.

**Author Summary:** Out of the three isoforms of JNKs (cJun N-terminal Kinases) in human (each with distinct cellular locations and functions), JNK1 plays role in starvation induced activation of autophagy. The role of JNK1 in autophagy modulation and dual role of autophagy in tumor cells makes JNK1 a promising anticancer drug target. Since JNKs are closely related to other members of MAPK (Mitogen-Activated Protein Kinases) family, it is difficult to design JNK selective inhibitors. Designing JNK isoformselective inhibitors are even more challenging as the ATP binding sites among all JNKs are highly conserved. Random forest classifier usually outperforms several other machine learning algorithms for classification and prediction tasks in diverse areas of research. In this work, we have used Random Forest algorithm for two purposes: (i) calculating variable importance measures to rank and select molecular features, and (ii) predicting JNK1 inhibitors regulating autophagy in cancer. We have used paDEL calculated molecular features of JNK1 bioactivity dataset from ChEMBL database to build classification models using random forest classifier. Our results show that by optimally selecting features from top 10% based on variable importance measure the classification accuracy is high, and the classification model proposed in this study can be integrated with drug design pipeline to virtually screen compound libraries for predicting JNK1 inhibitors.

## 1 Introduction

Two Nobel prizes in physiology or medicine, one for the discovery of lysosomes to Christian de Duve in 1974 and another for the discovery of mechanisms of autophagy to Yoshinori Yosumi in 2016, attracted many reseachers to work towards and uncover the fundamental physiological importance of autophagy for human health and diseases. Autophagy is a catabolic process by which heterogenic cellular materials are delivered to lysosomal digestion. Various diseases such as tumorigenesis, neurodegeneration, immune diseases as well as ageing are caused by chemical, genetic and age-driven changes in autophagic activity (Dikic and Elazar, 2018). Although the core molecular components involved in the execution of autophagy are well studied, there is limited information on how cellular signaling pathways, particularly kinases, regulate this complex process. The human kinome constitutes about 2% of all human genes, comprising around 538 eukaryotic protein kinase (ePK) genes which are subdivided into seven families of typical and seven families of atypical protein kinases, majority of which are serine/threonine protein kinases (STPKs). Kinases are integral to autophagy. Atg1, the first autophagy-related protein identified is a STPK, and is regulated by another STPK-mTOR. The role of many different kinases in regulation of various components/steps of autophagy is discussed in literature. mTOR, AMPK (AMP-activated protein kinase), Akt, MAPK (mitogen-activated protein kinase viz. ERK, p38 and JNKs) and PKC (protein kinase C) are other STPKs often deregulated in cancer and hence are important therapeutic targets (Sridharan et al., 2011).

JNKs (c-Jun N-terminal kinases) are also known as stress-activated protein kinases (SAPKs) as they are activated in response to inhibition of protein synthesis. The JNKs bind and phosphorylate the DNA binding protein c-Jun (a component of the AP-1 transcription complex which is an important regulator of gene expression) and increase its transcriptional activity. There are three isoforms of JNKs in mammals-JNK1, JNK2 and JNK3, each with distinct cellular locations and functions. In mammalian cells, the antiapoptotic protein, Bcl-2, binds to Beclin 1 during nonstarvation conditions and inhibits its autophagy function. Experimental studies show that JNK1 plays role in starvation induces phosphorylation of cellular Bcl-2 at residues T69, S70, and S87 of the nonstructured loop which causes dessociation of Bcl-2 from Beclin 1 and activation of autophagy. It is found that JNK1 but not JNK2 plays role in starvation induced activation of autophagy (Wei et al., 2008), while JNK3 is implicated in neuronal apoptosis (Xie et al., 1998). JNKs are closely related to other members of MAPK family such as the p38, MAP kinases and the ERK (extracellular-regulated kinase), hence it is difficult to design JNK-selective inhibitors. Designing JNK isoform-selective inhibitors are even more challenging as the ATP-binding sites among all JNKs are highly conserved (Koch et al., 2014).

As of February 2015, there are 33 approved kinase inhibitors most of which are launched for the treatment of cancer except tofacitinib (a JAK3 inhibitor) for the treatment of rheumatoid arthritis, sirolimus for organ rejection, fasudil for cerebral vasospasm, and nintedanib(a VEGFR2 inhibitor) for the treatment of Idiopathic pulmonary fibrosis, while more than 130 kinase inhibitors are reported to be in Phase-2/3 clinical trials. The design of selective kinase inhibitors is challenging because of structural similarity in the ATP binding site, which is the target of most of the approved and clinically advanced kinase inhibitors except rapalogs and trametinib. The poor selectivity of the kinase inhibitors within the kinome leads to undesirable side effects (Fabbro et al., 2015; Roskoski Jr, 2016). Several highly selective pan-JNK inhibitors have been characterized, and the first potent pan-JNKs inhibitor-SP600125 (Celgene) is still a common reference in JNK assay systems. Few chemical entities targeting JNKs– Bentamapimod (AS602801, PGL5001), CC-930 (tanzisertib, F2) and a peptidic inhibitor XG-102 (AM-111, O2) have been in clinical trials. Since JNK proteins may promote tumour development in a tissue‐ or cell-specific manner, designing JNK-isoform selective inhibitors (either ATP-competitive/non-competitive or inhibiting downstream cellular targets of a specific JNK protein in a tissue-specific manner) may be effective in checking specific tumour formation (Bubici and Papa, 2014). However, it is difficult to design isoform-specific JNK inhibitor, and such inhibitors are not yet commercially available (Gehringer et al., 2015). Crystal structures of JNK1 with its inhibitors have been reported from Protein Data Bank (PDB ids-3ELJ, 4AWI, 4L7F, 3PZE, 2NO3, 2GO1) and bioactive molecules of JNK1 are deposited in biological databases. Together, these may guide structure-based or ligand-based inhibitor design for JNK1 kinase. Compound classification and similarity search based approaches are two broad ligand-based approaches commonly applied for virtual screening of large compound library. Clustering, partitioning and machine learning-based approaches are well known compound classification based approaches, whereas Molecular graphs (2D) or conformations (3D) derived molecular fingerprints, and 3D pharmacophore models are few similarity search based approaches well adopted in ligand-based virtual screening (LBVS). Several machine learning-based classification algorithms are available such as Decision Tree (DT), Random Forest (RF), Support Vector Machine (SVM), Naive Bayes (NB), K-Nearest Neighbour (KNN) and Artificial Neural Network (ANN) (Lavecchia, 2015). These LBVS approaches can be applied irrespective of target information.

Detail studies of JNK signalling pathways in cancer to design and develop JNK inhibitors are reviewed in literature (Bubici and Papa, 2014; Messoussi et al., 2014). Yao et al., designed a selective inhibitor of JNK1, AV-7, and tested in *in vitro* studies (Yao et al., 2009). Kataria et al., studied inhibitor design against JNK1 through e-pharmacophore modeling docking and molecular dynamics simulations (Katari et al., 2016). To date, limited informations are available with focus on computational studies to design JNKs inhibitors, while it is diificult to find literatures on prediction of isoform-specific JNKs inhibitors using machine learning techniques. Hence, in this study we have used ligand-based machine learning approach using random forest to predict JNK1 inhibitor regulating autophagy in cancer.

## 2 Material and Methods

### 2.1 Data Set

The whole data set is downloaded from ChEMBL (Gaulton et al., 2012) using the following refining criteria: (1) only human JNK1 inhibition assay data based on enzyme or enzyme regulation are collected (2) duplicated compounds and compounds without detail assay value (IC50) are not considered. By applying these criteria, 1486 diverse compounds associated with JNK1 (target id CHEMBL2276) are selected for our study which have IC50 values ranging from 0.00024*μ*M to 667*μ*M. The 2-D structures of the compounds are converted into 3-D structures using CORINA (version 2.64) software. The molecules are saved in 3-D sdf format. Finally, 1198 compounds with calculated descriptors are selected in which 988 with IC50 values less than 10*μ*M are considered *active* and remaining 210 compounds with IC50 values ≥ 10*μ*M are considered *inactive*. For hit-to-lead activity of bioactives from biological databases, 10*μ*M cutoff value is considered to be reasonable starting point for most of the targets unless specifically specified in literature through experimentations.

### 2.2 Calculation and Selection of Molecular Descriptors

Prior to splitting the data set in training and test set, molecular descriptors of JNK1 bioactives are calculated using PaDEL descriptor software (Yap, 2011). Informative descriptors are retained and prioritized using Random Forest Variable Importance Measures (RFVIMs), while non-informative descriptors are removed from the final selected data sets.

#### 2.2.1 PaDEL-Descriptor

The software currently calculates 1875 descriptors (1444 1D, 2D descriptors and 431 3D descriptors) and 12 types of fingerprints. The descriptors and fingerprints are calculated using Chemistry Development Kit with few additional descriptors and fingerprints such as atom type electrotopological state descriptors, atom-based calculation of partition coefficient (Crippen’s logP) and Molecular Refractivity (MR), extended topochemical atom (ETA) descriptors, molecular linear free energy relation descriptors, ring counts, count of chemical substructures identified by Laggner, and binary fingerprints and count of chemical substructures identified by Klekota and Roth.

In present study, 1444 (1D and 2D) descriptors of JNK1 bioactives are calculated using PaDEL descriptor software (Yap, 2011) prior to splitting the data set in training and test set. 269 out of which 1444, 1-D and 2-D descriptors are finally selected, as descriptors with NA (not applicable) for most of the samples are removed. Finally, 1198 unique samples with known IC50 values *(988active* and 210 *inactive*) and 269 calculated features of each samples are used for model building using RF algorithm. Few calculated descriptors in this study are– Atom type electrotopological state (nHBint6), Molecular distance edge (MDEN-22), Atom type electrotopological state (SHBint6), Molecular distance edge (MDEN-12), Atom type electrotopological state (MaxHBint6), Atom type electrotopological state (minHBint6), Molecular linear free energy relation (MLFER A), Extended topochemical atom (ETA BetaP ns d), Autocorrelation (AATSC6c), Atom type electrotopological state (minssNH) etc.

#### 2.2.2 Variable Importance Measures (VIMs)

The permutation VIM and the Gini VIM are two different VIMs calculated for each predictor in standard Random Forest suggeted by Brieman. Permutation VIM is defined as the difference between the out-of-bag (oob) error resulting from a data set obtained through random permutation of the predictor of interest and the oob error resulting from the original data set. The oob error is expected to increase on permutation of an important predictor, which leads to high permutation VIM. The Gini VIM of a predictor of interest is the sum of the DGI (decrease of Gini impurity) criteria of the splits that are based on this predictor, scaled by the total number of trees in the forest. An important predictor is often selected for splitting and yields a high DGI when selected, leading to a high Gini VIM (Boulesteix et al., 2012). Here, VIMs of predictors are calculated to retain the informative descriptors and remove the non-informative descriptors using IncMSE (based on *mean square error*) and IncNodePurity (based on *node purity*) functions of Random Forest library in R package.

### 2.3 Machine Learning-based Compound Classification

Machine learning-based compound classification for activity prediction needs prior knowledge of annotated compounds with specific activity for designing a training set divided into *active* or *inactive* class. The training set is then analyzed to develop classifier. A classifier is a function that assigns a label or class to an unlabeled sample. In this study, a Random Forest (RF) classifier is developed to classify JNK1 bioactives as inhibitor or non-inhibitor. The complete data set of 1198 bioactive molecules selected from ChEMBL database are divided into five folds using random sampling technique. One fold is used as test set while four folds are used to train the model. The training and testing processes are repeated five times, each time with different fold as test set and remaining four folds to train the model, so that every molecule get trained and tested at least once. This method is known as five-fold cross validation using which the robustness of the trained model is checked.

#### 2.3.1 Random Forest

In RF (Breiman, 2001) algorithm, the unpruned classification or regression trees are generated based on random feature selection (Svetnik et al., 2003). RF builds and averages a large collection of mutually related trees. RF is significant modification of bagging or bootstrap aggregation which reduces the variance of an estimated prediction function. Following random sampling, when the training set for the present tree is drawn, about one-third of the data are left out of the sample. The use of out-of-bag (oob) sample is one of the important feature of RF which gives an ongoing unbiased estimate of the classification error as the number tree increases. oob error estimate is almost identical to the error obtained by n-fold cross-validation, which is being performed in parallel. Hence, the training can be terminated on stabilization of oob error (Friedman et al., 2001). Two main applications of RF technique are: (i) to assess and rank variables by their discriminative power, and (ii) to construct classifier for supervised learning problem. RF is an ensemble learning algorithm as predictions are made by aggregating majority vote or averaging the predictions of the ensemble. When the number of predictors is much larger than the number of observations, RF is capable of showing excellent performance. The application of RF as a standard data analysis tool in bioinformatics is reviewed by Boulesteix et al. (2012). Random Forest is implemented in a variety of packages including R.

## 3 Results

### 3.1 Prioritization of Molecular Descriptors

Random Forest automatically selects and ranks the important descriptors using IncNodePurity and IncMSE functions for VIM. The model uses regression for calculating VIM and shows out-of-bag (oob) error of 0.08675 (mean square error; MSE) which corresponds to 60.93% variance unexplained when number of trees (ntree) is 500 and the number of input variables to be used in each node (mtry) is 89 as shown in Table 1. The oob data (one-third of training data) is able to explain 39.07% variance of the training data. The model shows that the oob error remains almost constant after 300 trees are generated as shown in Fig.1. We have considered 500 trees to avoid over-fitting of the model.

**Table 1:** RF model to calculate VIM showing no. of trees, mean square error and %variance

| # Trees | MSE | %variance |
| --- | --- | --- |
| 100 | 0.08841 | 62.10 |
| 200 | 0.08725 | 61.28 |
| 300 | 0.08734 | 61.34 |
| 400 | 0.08709 | 61.16 |
| 500 | 0.08675 | 60.93 |

**Figure 1:**
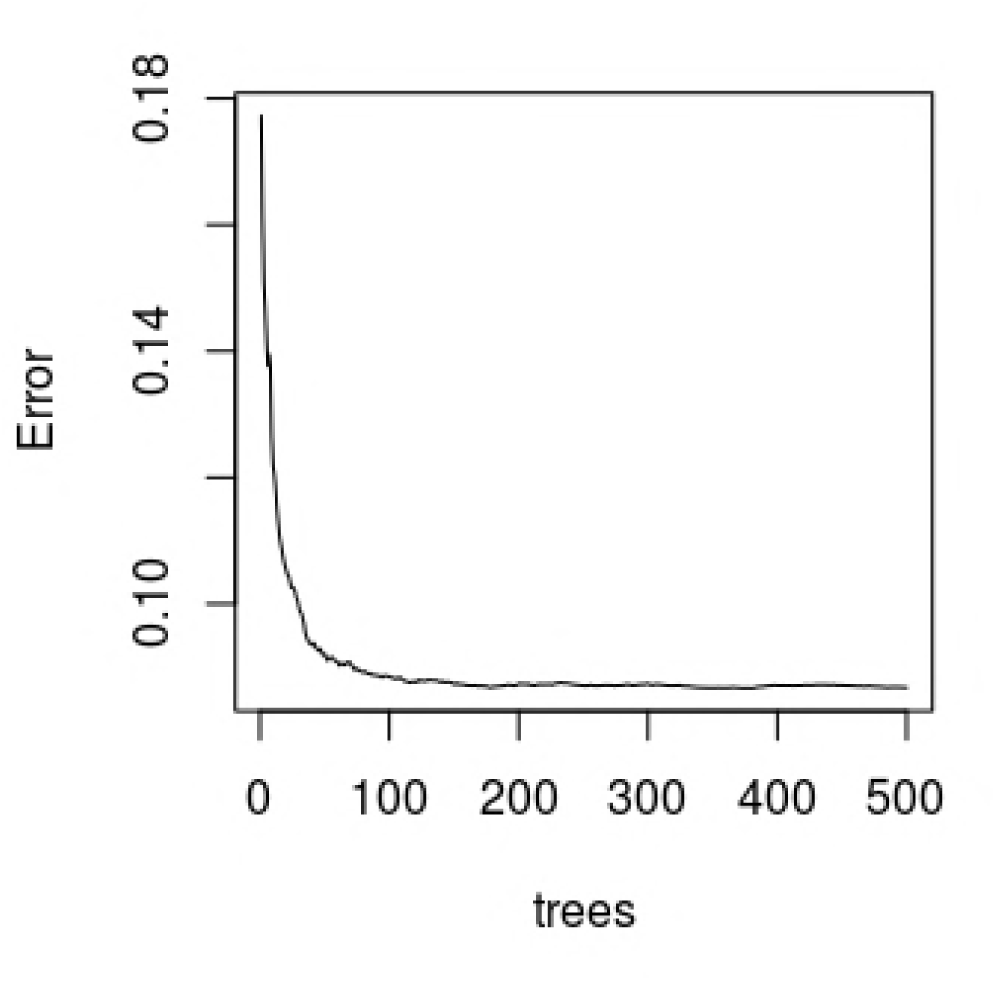
RF model to calculate VIM (showing no. of trees and out-of-bag error)

Out of 269 calculated descriptors, top 10% (27) descriptors are selected for building classification model using RF classifier. Top 10 descriptors with their IncMSE measures are shown in table 2, whereas top 30 descriptors with their IncMSE measures are plotted as shown in Fig. 2.

**Table 2:** Relative importance of top 10 descriptors using IncMSE function of RF-VIM

| Rank | Descriptors | IncMSE |
| --- | --- | --- |
| 1 | ATSC0c | 16.8522455505 |
| 2 | AATSC1v | 12.6246738825 |
| 3 | AATS7p | 12.4114747628 |
| 4 | ATSC3s | 11.6990337935 |
| 5 | AATSC3s | 11.4763612558 |
| 6 | AATSC0c | 10.8516271753 |
| 7 | MATS3m | 10.1128028665 |
| 8 | ATSC1v | 9.0966608707 |
| 9 | AATS4i | 9.0962395998 |
| 10 | AATSC8p | 8.6634381244 |

**Figure 2:**
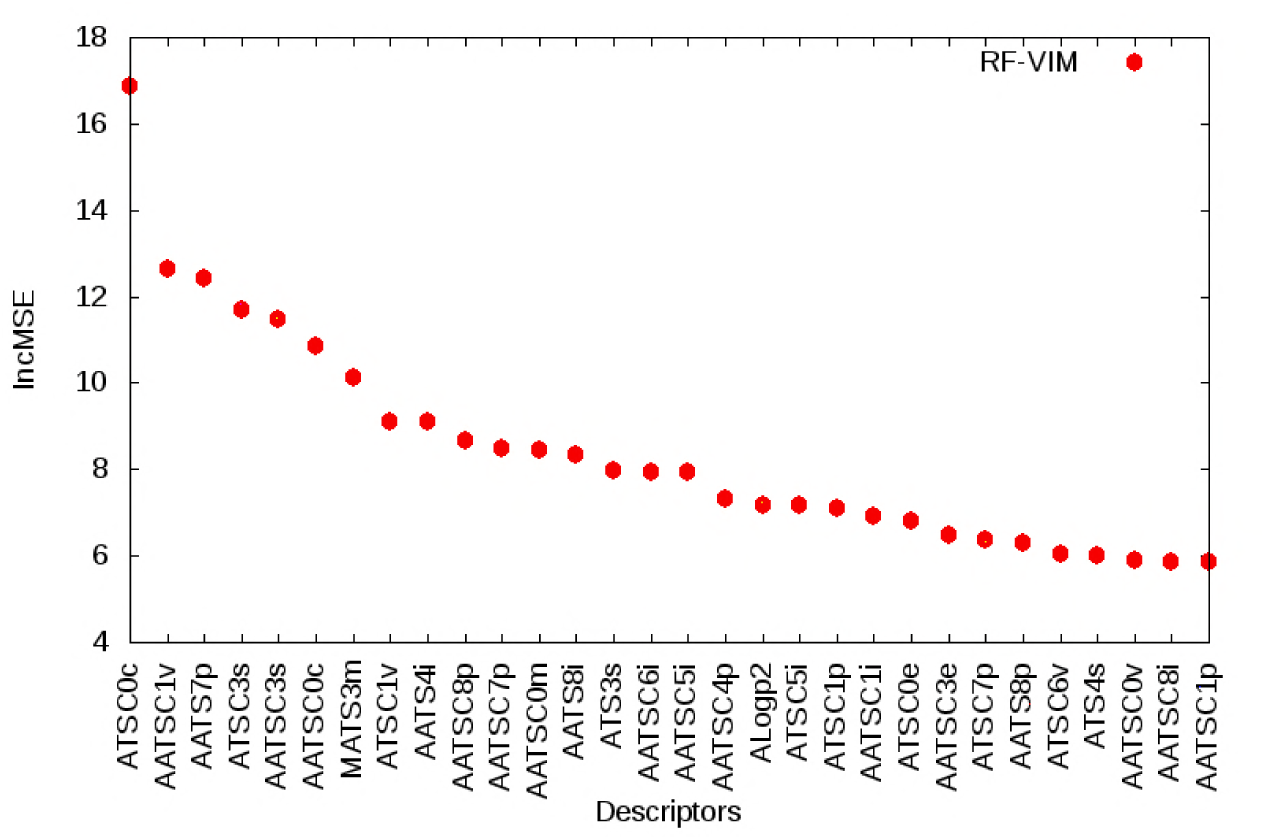
Relative importance of top 30 variables using IncMSE function of RF.

Top 10 descriptors with their IncNodePurity measures are shown in table 3, whereas top 30 descriptors with their IncNodePurity measures are plotted as shown in Fig. 3.

**Table 3:** Importance of top 10 variables using IncNodePurity function of RF-VIM

| Rank | Descriptors | IncNodePurity |
| --- | --- | --- |
| 1 | ATSC0c | 5.1225495212 |
| 2 | AATS7p | 3.0129326771 |
| 3 | AATSC0c | 2.3678963858 |
| 4 | AATSC3s | 2.0567042176 |
| 5 | AATSC1v | 1.9148403892 |
| 6 | ATSC3s | 1.7456336954 |
| 7 | AATSC7p | 1.5572340296 |
| 8 | AATSC6i | 1.4527806122 |
| 9 | ATSC1v | 1.4285816216 |
| 10 | AATS4i | 1.3773576137 |

**Figure 3:**
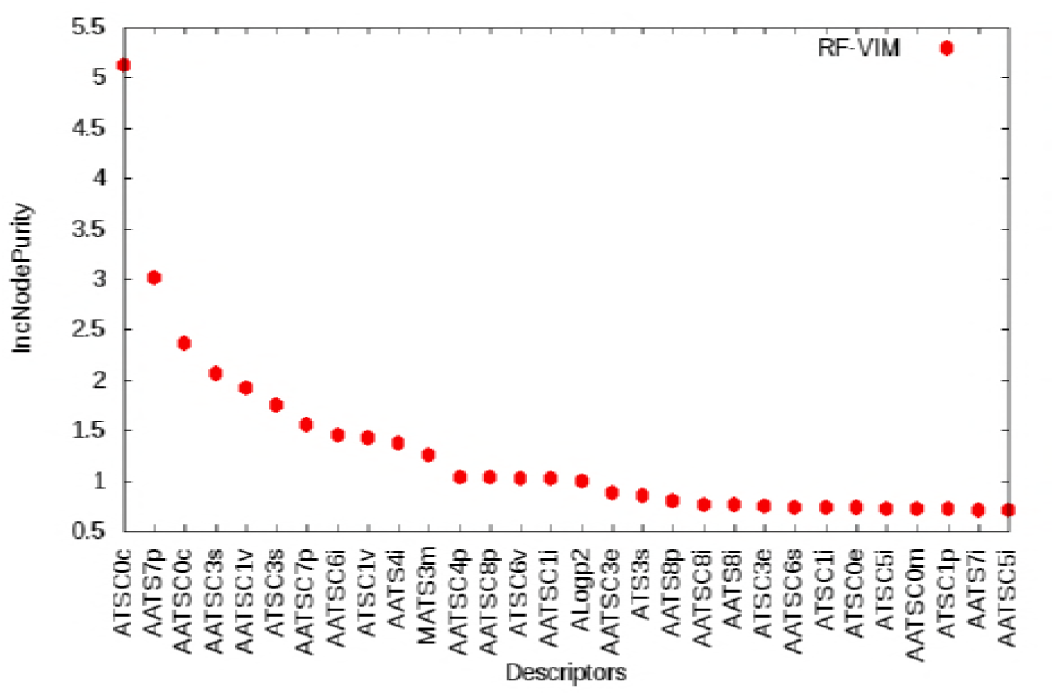
Relative importance of top 30 variables using IncNodePurity function of RF.

Using IncNodePurity (>0.720) and IncMSE (>5.9) of RF-VIM, top 10% (27) descriptors are extracted using both functions. Finally, 22 common variables are selected for building classification models using RF classifier.

### 3.2 Performance Evaluation and Cross-Validation of the Model

Various performance evaluation metrics viz. Accuracy, Specificity, Recall (Sensitivity) and Precision (Positive Prediction Value) are calculated using confusion matrix as shown in Table 4. These metrics are calculated using number of inhibitors or True Positives (TP), non-inhibitors or True Negatives (TN), non-inhibitors predicted as inhibitors or False Positives (FP), and inhibitors predicted as non-ihibitors or False Negatives (FN). Cross-validation is performed to validate the accuracy and robustness of the prediction model. In our experiment, 5-fold cross-validation is used to validate the generated models using RF (default setting).

**Table 4:** Confusion matrix of the classifiers based on top 10% of selected descriptors

| Classifier | Folds | 1 | 2 | 3 | 4 | 5 | Sum |
| --- | --- | --- | --- | --- | --- | --- | --- |
| RF | TP | 188 | 201 | 189 | 184 | 186 | 948 |
|  | TN | 18 | 17 | 12 | 16 | 21 | 84 |
|  | FP | 25 | 15 | 33 | 30 | 23 | 126 |
|  | FN | 8 | 6 | 5 | 9 | 9 | 37 |

By default, RF picks up 2/3rd data for training and 1/3rd data for testing for regression and almost 70% data for training and 30% for testing during classification. Since it randomizes the variable selection during each tree split, it is not prone to overfit unlike other models. Out-of-bag error calculated during model training is an indicator of test set performance. However, to ensures that all samples appear in the training and test sets so that 100% of the data gets used at some point for training and for testing, cross validation is crucial for model evaluation and comparison with other models for the same data set. Hence, 5-fold cross-validation is used for model evaluation.

The data set of 1198 molecules which include *988active* (IC50 values <10*μ*M) and 210 *inactive* (IC50 values ≥ 10*μ*M) are randomly sampled followed by five-fold cross-validation. The complete data set is split into five parts so that four parts (959) are considered for training and remaining one part (239) for testing. The process is iterated five times so that each time different set of molecules are considered for testing. Then the models are developed with classifier using RF to classify the molecules as inhibitors or non-inhibitors. For evaluation of the models, different statistical measures such as Accuracy, Specificity, Sensitivity (Recall) and Precision are calculated. When the classifiers are based on top 10% of selected descriptors, RF model shows accuracy of 86.36%, precision of 88.27%, specificity 40.00% and sensitivity 96.24%, as is shown in table 4 & 5.

**Table 5:** Performance evaluation metrics of classifiers based on top 10% of selected descriptors

| Classifiers | Accuracy (%) | Specificity (%) | Sensitivity (Recall) | Precision (%) |
| --- | --- | --- | --- | --- |
| RF | 86.36 | 40.00 | 96.24 | 88.27 |

A ROC (Receiver Operating Characteristic) curve plots the true positive rate (sensitivity) against the false positive rate (1 — *specificity*) for all possible cutoff values. ROC curve for the RF classification model which is based on top 10% descriptors is shown in Fig.4.

**Figure 4:**
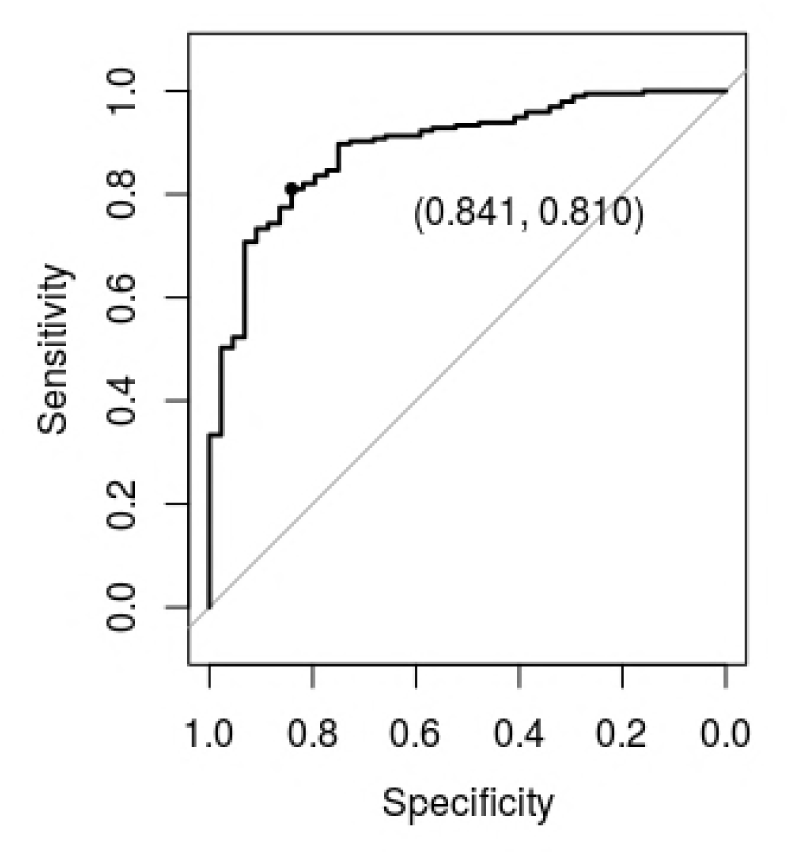
ROC curve of RF classifier based on common out of top 10% descriptors ranked by RF-VIM.

Classification model based on a set of 22 common molecular descriptors out of top 27 ranked by both incMSE and incNodePurity functions of RF-VIM achieves accuracy 86.36% and area under ROC curve (AUC) 0.8914 respectively using RF classifier.

## 4 Discussion

In this study, we have used machine learning to prioritize molecular descriptors, and based on which *in silico* classification models have been developed to classify a data set of 1198 bioactive molecules of JNK1 from ChEMBL database as inhibitor or inhibitor. 1444 (1D and 2D) molecular descriptors are calculated using PaDEL descriptor software, out of which 269 descriptors are retained after removing the descriptors with non-informative values (NA) for most of the compounds, and then ranked using IncMSE and IncNodepurity functions of Random Forest Variable Importance Measure (RF-VIM). As the out-of-bag error remains constant after 300 trees, this would be an optimal choice for model building for calculating RF-VIM. As no single descriptor is enough to discriminate inhibitors from non-inhibitors, a set of 22 descriptors out of top ranked 10% of 269 (27) descriptors are prioritized and selected to build the classification models. The data set of 1198 molecules which include *988active* (IC50 values <10*μ*M) and 210 *inactive* (IC50 values ≥ 10*μ*M) are randomly sampled followed by five-fold cross-validation. The complete data set is split into five parts so that four parts (959) are considered for training and remaining one part (239) for testing. The process is iterated five times so that each time different set of molecules are considered for testing. Then the classification models are developed using RF to classify the molecules as inhibitors or non-inhibitors. For evaluation of the models, different statistical measures such as Accuracy, Specificity, Sensitivity (Recall) and Precision are calculated. When the classifiers are based on top 10% of selected descriptors, RF model shows accuracy of 86.36%, precision of 88.27%, specificity 40.00% and sensitivity 96.24%.

## 5 Conclusion

Though virtual screening (VS) methods (either structure or ligand-based) cannot be solely applied to design a new drug, yet these methods when applied in the initial stages of drug development processes, dramatically reduce the time and cost investment to complete a drug development cycle. Machine learning-based compound classification is one of the ligand-based virtual screening (LBVS) method which is studied here to develop classification models to classify JNK1 bioactives from ChEMBL database as inhibitor or non-inhibitor. We have developed machine learning-based classification models using RF to predict JNK1 inhibitors. JNK1, a serine-threonine kinase has implications in several diseases including cancer. Based on the existing literature it is found that JNK1 has dual role in cancer through the process of autophagy. In mammalian cells, the antiapoptotic protein, Bcl-2 binds to Beclin 1 during nonstarvation conditions and inhibits its autophagy function. Experimental studies show that starvation induces phosphorylation of cellular Bcl-2 by JNK1 which causes dessociation of Bcl-2 from Beclin 1 and activation of autophagy. It is also found that JNK1 but not JNK2 plays role in starvation induced activation of autophagy (Wei et al., 2008), while JNK3 is implicated in neuronal apoptosis (Xie et al., 1998). Here, we have used Random Forest algorithm for two purposes – to measure variable importance for optimal selection of molecular descriptors and as a classifier. It is found that by selecting top 10% of descriptors ranked by Random Forest Variable Importance Measure, RF classifier shows accuracy of 86.36%, precision of 96.24% and area under ROC curve (AUC)– 0.8914. The model based on RF classifier in this study can be used for virtual high throughput screening of large compound libraries of JNK1 bioactives. The limitations of this study include choice of data sets (which should ideally be stage-specific for the study of autophagy regulating kinases (as autophagy acts as tumor-suppressor in the initial stage of tumor development and as a tumor-promoter in well settled tumor cells to endure stressful conditions), the choice of descriptors, prediction performance and the comparision with other classifiers viz. Support Vector Machine, Naive Bayes, Decision Tree. Including more autophagy regulating kinases as targets to build a common model to screen large compound library for autophagy regulating kinases is one of our future directions of work.

